# Early split between African and European populations of *Drosophila melanogaster*

**DOI:** 10.1101/340422

**Authors:** Adamandia Kapopoulou, Martin Kapun, Pavlos Pavlidis, Bjorn Pieper, Ricardo Wilches, Wolfgang Stephan, Stefan Laurent

## Abstract

Natural populations of the fruit fly *Drosophila melanogaster* have been used extensively as a model system to investigate the effect of neutral and selective processes on genetic variation. The species expanded outside its Afrotropical ancestral range during the last glacial period and numerous studies have focused on identifying molecular adaptations associated with the colonization of northern habitats. The sequencing of many genomes from African and non-African natural populations has facilitated the analysis of the interplay between adaptive and demographic processes. However, most of the non-African sequenced material has been sampled from American and Australian populations that have been introduced within the last hundred years following recent human dispersal and are also affected by recent genetic admixture with African populations. Northern European populations, at the contrary, are expected to be older and less affected by complex admixture patterns and are therefore more appropriate to investigate neutral and adaptive processes. Here we present a new dataset consisting of 14 fully sequenced haploid genomes sampled from a natural population in Umeå, Sweden. We co-analyzed this new data with an African population to compare the likelihood of several competing demographic scenarios for European and African populations. We show that allowing for gene flow between populations in neutral demographic models leads to a significantly better fit to the data and strongly affects estimates of the divergence time and of the size of the bottleneck in the European population. Our results indicate that the time of divergence between cosmopolitan and ancestral populations is 30,000 years older than reported by previous studies.

## Introduction

*Drosophila melanogaster* originated in sub-Saharan Africa where it diverged from its sister species *Drosophila simulans* approximately 2.3 million years ago (David & Capy, 1988). Accordingly, South and East African populations display genetic diversity patterns closer to mutation-drift expectations compared to western African and non-African populations, providing further evidence that this geographic area represents the ancestral range of the species (David & Capy, 1988; Haddrill, Thornton, Charlesworth, & Andolfatto, 2005; Veuille, Baudry, Cobb, Derome, & Gravot, 2004). Previous genetic analyses of European and Asian samples indicated that non-African populations started expanding beyond their ancestral range around 13,000 years ago, eventually colonizing large areas in Europe and Asia (Laurent, Werzner, Excoffier, & Stephan, 2011; Li & Stephan, 2006). By contrast, the introduction of the species in the Americas and Australia is very recent (within a couple of hundred years ago) and has been well documented by early entomologists (reviewed in Keller, 2007). Interestingly, demographic analyses of a North-American and Australian populations revealed significant African ancestry (between 15 and 40%) in a dominantly European background (Bergland, Tobler, Gonzalez, Schmidt, & Petrov, 2016; Caracristi & Schlotterer, 2003; Duchen, Zivkovic, Hutter, Stephan, & Laurent, 2013; Kao, Zubair, Salomon, Nuzhdin, & Campo, 2015).

Natural populations of *D. melanogaster* have also been used extensively to study the effect of positive and negative selection on functional and linked neutral variants (reviewed in Casillas and Barbadilla, 2017; Charlesworth, 2012; Sella, Petrov, Przeworski, & Andolfatto, 2009), to obtain estimates for the rate of adaptive events and the magnitude of the fitness effects of beneficial mutations, and to identify genes displaying molecular signatures of hitchhiking events. However, these studies also emphasized the necessity - and difficulty - to jointly consider the effects of positive selection, background selection, and demographic processes (Elyashiv et al., 2016). Studying the joint effect of neutral and selective forces on genetic variation has been facilitated by the recent sequencing of large numbers of complete genomes from natural populations (Grenier et al., 2015; Lack et al., 2015; Lack, Lange, Tang, Corbett-Detig, & Pool, 2016; Langley et al., 2012; Mackay et al., 2012; Pool et al., 2012). However, most non-African full-genome datasets have been obtained from new world populations implying that analyses of this material must account for recent genome-wide admixture. A small number of European and Asian inbred lines have been sequenced recently (Grenier et al., 2015), but this data retains large regions of residual heterozygosity (>500kb), which represents a challenge for population genomic studies (Langley, Crepeau, Cardeno, Corbett-Detig, & Stevens, 2011), because the nature of the sequenced biological material (inbred lines) does not allow obtaining phased data.

Here, we followed a protocol proposed by Langley et al. (2011) and sequenced haploid genomes of 14 *Drosophila melanogaster* embryos obtained from a Swedish population. We describe patterns of genetic diversity and compare these to previously available data from a Zambian population located in the ancestral range of the species. We use this new dataset to re-visit different competing hypotheses concerning the demographic history of European populations. We show that accounting for historical gene flow in demographic models of European and African populations improves the fit to the data compared to previously published models and that, as a consequence, the estimate for the divergence time between African and non-African gene pools is older than previously reported.

## Materials and Methods

### Data collection

A total of 96 inseminated female *D. melanogaster* were sampled from Umeå in northern Sweden in August 2012. Then, full-sibling mating was performed for 10 generations, which yielded 80 inbred lines. Out of these, 14 lines were randomly selected from which haploid embryos were generated following the protocol described by (Langley et al., 2011). Standard genomic libraries were constructed using up to 10 μg (~200 ng/μl) of DNA. Library construction and sequencing of one haploid embryo for each of the 14 haploid-embryo lines were carried out on an Illumina HiSeq 2000 sequencer at GATC Biotech (Konstanz, Germany). In addition to the newly established and sequenced inbred lines from Umeå/Sweden, we randomly chose 10 lines not carrying any inversions from the DPGP3 dataset. They were collected in Siavonga/Zambia in July 2010 and sequenced as haploid embryos similar to our data. Since four of the Swedish lines carried the chromosomal inversion *In*(*2L*)*t*, we additionally chose four lines at random from Zambian lines that also carried *In*(*2L*)*t* to match the number and distribution of inversion karyotypes in our Swedish dataset (see Table S1).

### Mapping pipeline

Prior to mapping, we tested raw read libraries in FASTQ format for base quality, residual sequencing adapter sequences and other overrepresented sequences with FASTQC (v0.10.1; http://www.bioinformatics.babraham.ac.uk/projects/fastqc/). We trimmed both the 5’ and 3’ end of each read for a minimum base quality ≥ 18 and only retained reads with a minimum sequence length ≥ 75bp using cutadapt (v 1.8.3 Martin, 2011). We used bbmap (v 35.50 Bushnell, 2017) with standard parameters to map intact read pairs, where both reads fulfilled all quality criteria, against a compound reference consisting of the genomes from *D. melanogaster* (v6.12) and genomes from other common pro- and eukaryotic symbionts including *Saccharomyces cerevisiae* (GCF_000146045.2), *Wolbachia pipientis* (NC_002978.6), *Pseudomonas entomophila* (NC_008027.1), *Commensalibacter intestini* (NZ_AGFR00000000.1), *Acetobacterpomorum* (NZ_AEUP00000000.1), *Gluconobacter morbifer* (NZ_AGQV00000000.1), *Providencia burhodogranariea* (NZ_AKKL00000000.1), *Providencia alcalifaciens* (NZ_AKKM01000049.1), *Providencia rettgeri* (NZ_AJSB00000000.1), *Enterococcus faecalis* (NC_004668.1), *Lactobacillus brevis* (NC_008497.1), and *Lactobacillus plantarum* (NC_004567.2) to avoid paralogous mapping of reads belonging to different species. We further filtered for mapped reads with mapping qualities ≥ 20, removed duplicate reads with Picard (v2.17.6; http://picard.sourceforge.net) and re-aligned sequences flanking insertions-deletions (indels) with GATK (v3.4-46 McKenna et al, 2010)

### Quality control

Since all libraries were constructed from haploid embryos, we assumed that polymorphisms within a library represent either (1) sequencing or (2) mapping errors. Accordingly, we expected to find erroneous alleles only at very low frequencies in each dataset. Alternatively, any problem during the construction of haploid embryos would lead to diploid sequences that result in residual heterozygosity characterized by an excess of polymorphisms with frequencies close to 0.5 in the affected library. To test for these hypotheses, we investigated the distribution of minor – putatively erroneous – allele frequencies for each library separately. In addition, we divided the number of erroneous alleles by the total coverage at variant and invariant positions to calculate library-specific error-rates.

### Variant calling

We identified single nucleotide polymorphisms (SNPs) based on a combination of stringent heuristic criteria to exclude sequencing and mapping errors in each of the Swedish and Zambian datasets using custom software: For each library, we excluded polymorphic positions with minor frequencies > 0.1. In all other cases, we considered the major allele as the correct allelic state for a given individual. To avoid erroneous SNPs due to inflated sampling error at low-coverage sites or due to paralogous alleles at sites with excessive coverage from mapping errors, we only considered positions with more than 10-fold and less than 200-fold coverage. We further ignored positions where less than 14 of the 28 samples (14 Swedish and 14 Zambian) fulfilled the above-mentioned quality criteria. At last, we refined the SNP dataset by excluding SNPs located either within known transposable elements (TE) based on the *D. melanogaster* reference genome (v.6.12) or within a 5 base-pairs distance to indel polymorphisms supported by 10 reads across all samples. Finally, the same set of filters was applied to each other non-polymorphic chromosomal position in the data. This allowed us to obtain the total number of monomorphic sites in our dataset, which is needed for demographic inference (Laurent et al., 2016).

### Bioinformatic karyotyping

Following the approach in Kapun, van Schalkwyk, McAllister, Flatt, and Schlotterer (2014), we used a panel of karyotype-specific marker SNPs that are diagnostic for seven chromosomal inversions *(In(2L)t, In(2R)NS, In(3L)P, In(3R)C, In(3R)K, In(3R)Mo* and *In(3R)Payne)* to karyotype all Swedish samples based on presence or absence of alleles which are in tight linkage with the corresponding inversion. We further used the same method to confirm the inversion status in previously karyotyped samples from Zambia. We only considered a sample to be positive for an inversion if it carried ≥ 95% of all alleles that are specific to the corresponding inversion.

### Principal Component Analysis

Principal component analyses were conducted with the “auto_SVD” function from the R package *bigsnpr* (Prive, Aschard, Ziyatdinov, & Blum, 2017). This algorithm uses clumping instead of pruning to thin SNPs based on linkage disequilibrium, removes SNPs in long-range LD regions, and uses the thinned data to perform dimensionality reduction by singular value decomposition. Analyses were conducted on the full data and on each chromosomal arm separately.

### Demographic analyses

For demographic inference, we used SNPs from all neutral introns (smaller than 65bp, bases from the 8^th^ to the 30^th^ position, described as the most appropriate sites to be used for such analyses in Parsch, Novozhilov, Saminadin-Peter, Wong, and Andolfatto (2010) together with 4-fold degenerate sites present in chromosomes 2R, 3L, 3R, and X. The latter SNP class was obtained following Grenier et al. (2015). Autosomal and X-linked data were treated separately. All genomic regions spanned by common inversions were excluded from the analyses (as defined by coordinates of inversion breakpoints obtained from Corbett-Detig and Hartl (2012)). Additionally, long runs of Identity-By-Descent were masked from the African lines using a perl script available from the DPGP website (http://www.johnpool.net/genomes.html). Genomic regions that were identified as of European ancestry in the DPGP2 and DPGP3 dataset were not masked, because our demographic analyses were intended to evaluate the possibility of gene flow between the two populations. All coordinates were transformed to *D. melanogaster* genome reference version 6 using an in-house python script. In total, 390,852bp (42,306 SNPs) were used for the three autosomes together and 183,502bp (27,972 SNPs) for the chromosome X. We used the software *dadi* (Gutenkunst, Hernandez, Williamson, & Bustamante, 2009) to test four different demographic scenarios. In all models, the ancestral African population experienced a stepwise expansion at time T_exp_. After the expansion, (forward in time) the European population splits from the African population at time T_split_. Immediately after the split, the size of the new European population is instantaneously reduced to a population size Nbot, whereas the size of the African population does not change. After the bottleneck, the European population is allowed to recover exponentially until it reaches its current size Neu. The four scenarios differ in the modeling of migration following the population split. Model 1 (NOMIG) does not implement gene flow and is therefore similar to previously published models (Duchen et al., 2013; Laurent et al., 2011; Li & Stephan, 2006). Model 2 (SYMIG) implements symmetrical migration between the populations, starting immediately after the split and lasting until the end of the simulation (present). Model 3 (ASYMIG) is similar to model 2 but allows for asymmetrical migration rates. Finally, Model 4 (RASYMIG) is similar to model 3 except that asymmetrical migration only starts at time Tmig. These four models have six, seven, eight, and nine parameters, respectively. For every scenario, at least 10 independent runs with different initial parameters values were performed and the run achieving the highest likelihood was kept for parameter estimation and model choice. Model choice was done by comparing the Akaike information criteria (AIC) between models (Akaike, 1974). Confidence Intervals (CI) were calculated using the following procedure: First, 150 datasets were simulated using the best demographic model. These simulations were treated as pseudo-observed data and used to re-estimate demographic parameters under the best model. The set of 150 estimates for each demographic parameter was then used to construct the confidence intervals. Because the re-estimated parameters are not normally distributed, confidence intervals were calculated as the 2.5-97.5% percentiles (see Table 2). Nucleotide diversity, Tajima’s D, Fst, and the observed 1D and 2D site frequency spectra presented in Figure S3 were calculated with built-in functions implemented in *dadi*.

Past changes in population size were inferred with the program *MSMC* (Schiffels & Durbin, 2014). The analysis was performed on 20 line-pairs randomly drawn from the Swedish and Zambian populations, respectively, and on 40 pairs consisting of a single line drawn from each population at random. All available SNPs from chromosomes 2R, 3R, and 3L were used for this analysis. The software was invoked with the following options: *msmc -i 30-t 8 -p “20*1 +30*2*. The scaled times and the coalescence rates estimated by *MSMC* were converted to generations and Ne, respectively, using a per base-pair mutation rate of 1.3e-9 per bp/per generation (Laurent et al., 2011). This conversion of time-specific coalescence rates into N_e_ values is based on the assumption that the corresponding sample has been collected from a panmictic population. Indeed, it has been shown that in structured populations instantaneous coalescence rates in samples of size two can vary through time in the absence of any variation in N_e_ (Chikhi et al., 2018; Mazet, Rodriguez, Grusea, Boitard, & Chikhi, 2016). Therefore, Mazet et al. (2016) proposed to refer to N_e_ values estimated by *MSMC* (or any equivalent method) as inverse instantaneous coalescence rates (IICR). We adopt this terminology in this study and used the python script *estimIICR.py* (https://github.com/willyrv/IICREstimator) published by (Chikhi et al., 2018) to calculate the IICR profiles expected under the best demographic model inferred by *dadi* with autosomal data. This required to reformulate the model into the following *ms* command; *ms 2 1000000 -T-L -12 2 0 -m 1 2 10.0815052248 -m 2 1 2.41228728 -n 2 0.2064966823 -g 2 91.3987189817 -ej 0.0234640378 2 1 -en 0.03305336 1 0.4436970874* (Hudson, 2002).

## Results

### Summary statistics of mapping and patterns of missing data

Our sequencing effort of 14 Swedish lines yielded homogenous average coverage across all autosomal arms ranging from 53.3x on *2R* to 57.2x on *2L*. In contrast, we observed a slightly higher coverage on the *X* chromosome (63.7x). These patterns were consistent with the data from the Zambian lines, where we also found a slight coverage excess on the X. We, however, identified pronounced variation in library-specific read-depth, ranging from 19.4x in SU93n to 87.7x in SU02n for the 14 Swedish and to a lesser extent also in the Zambian lines, which ranged from 26.9x in ZI200 to 38.7x in ZI472 (Figure S1). We found no evidence for residual heterozygosity, which confirms that all sequenced libraries were based on haploid genomes only and are thus fully phased (see Figure S2). Furthermore, we observed that errors occurred at very low frequencies corresponding to an average error rate of 0.365% in the Swedish and 0.348% in the Zambian libraries. We observed very similar patterns of missing data between the Swedish and Zambian datasets after applying identical sets of quality filtering (Figure S3), with the exception of the Swedish line SU81n that contains 30% missing data.

### Patterns of genetic variation in the Swedish sample

Previous studies based on a smaller number of loci showed that European flies derived from an ancestral sub-Saharan population from which they diverged at the end of the last glacial maximum (Stephan & Li, 2007) and that the colonization was associated with a founder event during which European flies were subject to high genetic drift (Li & Stephan, 2006; Thornton & Andolfatto, 2006). This scenario predicts observable genetic differences between Swedish and Zambian flies as well as a lower amount of diversity in the former due to the population size bottleneck associated with the founding event. We used PCA analysis to explore whether these expectations were also observed in our new genome-wide diversity data (Figure 1). This analysis showed that the first principal component always clustered European and African samples separately and that Swedish lines consistently displayed smaller dispersion along the second principal component, reflecting lower diversity compared to the Zambian sample (Table 1, McVean, 2009). One important exception to this general pattern was observed on chromosome *2L*. In addition to the population specific clustering on PC1, we identified an equally strong clustering on PC2 that was perfectly consistent with the presence or absence of the known chromosomal inversion *In*(*2L*)*t*, whose occurrence in Sweden is here reported for the first time (Table S1). The effect of *In*(*2L*)*t* on population genetic structure has already been described in the DPGP3 dataset and, interestingly, has also been shown to extend beyond the chromosomal breakpoints of the inversion, which could reflect the effect of historical positive selection on the inverted arrangement (Corbett-Detig & Hartl, 2012).

**Figure 1:**
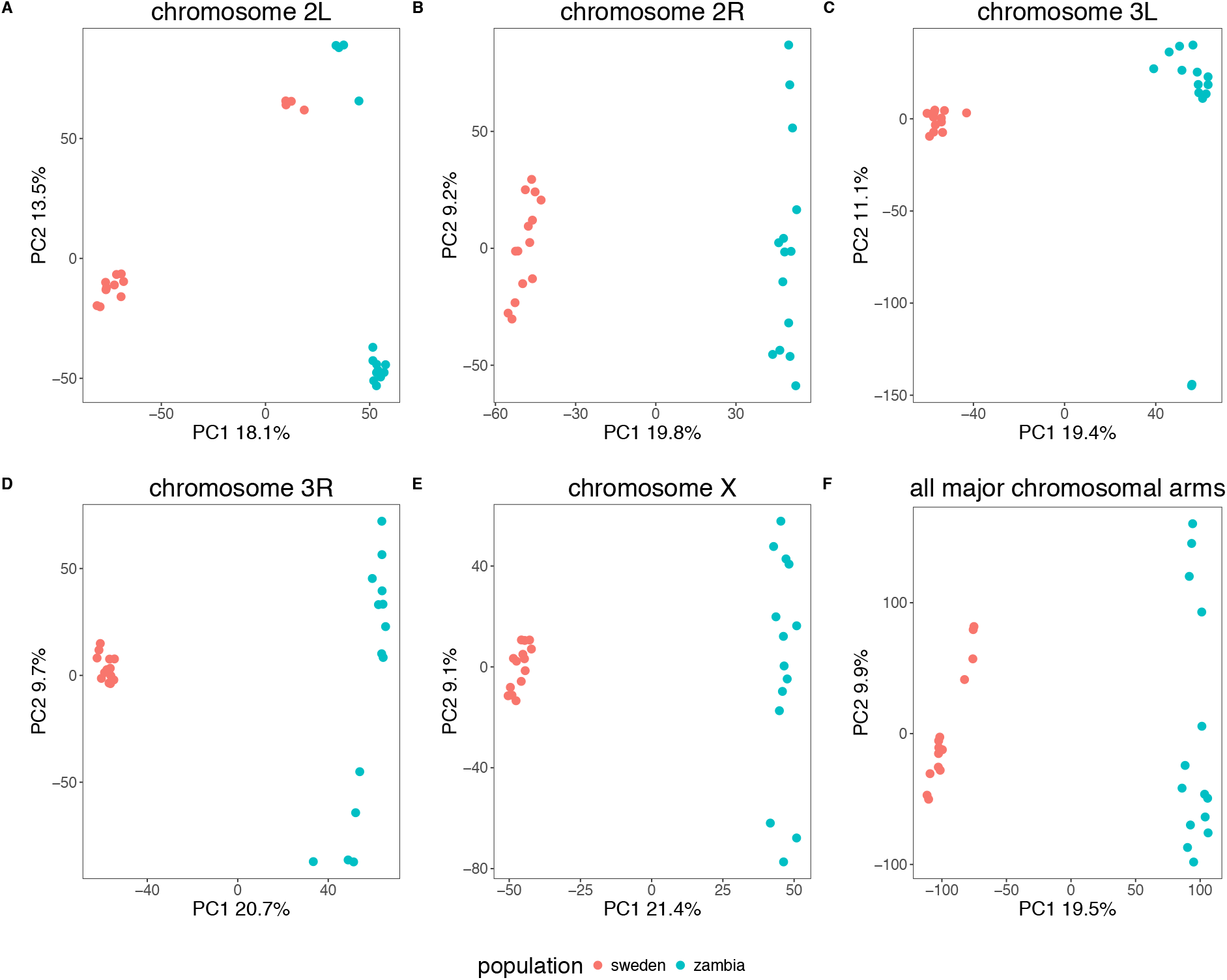
PCA results. Results are presented for each major chromosomal arm separately and for all chromosomes together. Only the first two components are shown. Individuals tend to cluster according to their sampling location except for chromosome 2L, for which flies carrying the inverted variant of the inversion *In*(*2L*)*t* cluster together regardless of their geographical origin.

**Table 1.** Summary statistics of genetic diversity measured on our neutral dataset (i.e. introns smaller than 65bp, and 4-fold degenerated third codon positions). All known inversions have been removed as well as chromosome 2L. All statistics have been calculated with *dadi* on the same site frequency spectra used for demographic inference.

|  | Umeå (Sweden) |  | Siavonga (Zambia) |  |
| --- | --- | --- | --- | --- |
|  | Autosomes | X | Autosomes | X |
| **$\theta_w$ (per bp)** | 0.007 | 0.005 | 0.013 | 0.014 |
| **$\theta_\pi$ (per bp)** | 0.008 | 0.005 | 0.012 | 0.013 |
| **Tajima's D** | 0.16 | 0.32 | -0.36 | -0.475 |
| **$F_{ST}$ (Umeå - Siavonga)** | 0.2 | 0.26 |  |  |

### Demographic modeling

To test whether migration represented an important evolutionary force after the split between the European and African populations, we designed four demographic models recapitulating the main assumptions about the possible role of migration in this system (Figure 2, see Materials and Methods for a description of the models). Model choice and parameter estimation were conducted using the software *dadi* 1.7.0 with a neutral subset of the data (see Materials and Methods). Population genetic statistics of the observed data (Table 1, Figure S4) were in line with values reported by previous studies based on smaller numbers of loci (Ometto, Glinka, De Lorenzo, & Stephan, 2005). Our demographic analyses showed that models including migration provided a better fit to the neutral data compared to the model without migration for both the autosomal and the X-linked dataset (Figure 2). For the autosomal data, the best fit was provided by model “ASYMIG” (ongoing asymmetrical migration, Figure 2). Under this model, divergence between the Zambian and Swedish samples for the autosomal data occurred 43,540 years ago (assuming 10 generations per year) and was followed by ongoing asymmetrical migration with the migration rate from Sweden to Zambia (M_SZ_=2Nm_SZ_=2.24) being larger than from Zambia to Sweden (M_ZS_=0.53). As expected, including gene flow into the models yielded older estimates for the age of the population split (Table 2, Table S2). We accordingly report here older divergence time than previous studies who did not take migration into account (Duchen et al., 2013; Laurent et al., 2011; Li & Stephan, 2006). For the X-chromosomal data, the best model was “SYMIG” (ongoing symmetrical migration). Under this model, divergence time was estimated to be 25,999 years with an ongoing symmetrical migration rate of 1.23 (number of genomes migrating per generation). X-chromosome modeling also confirmed the stronger estimated bottleneck for the X versus autosomes (Hutter, Li, Beisswanger, De Lorenzo, & Stephan, 2007; Laurent et al., 2011). The comparison between observed data and predictions of the best models showed that our modeling approach yielded a good absolute fit to the data (Figure S5).

**Figure 2:**
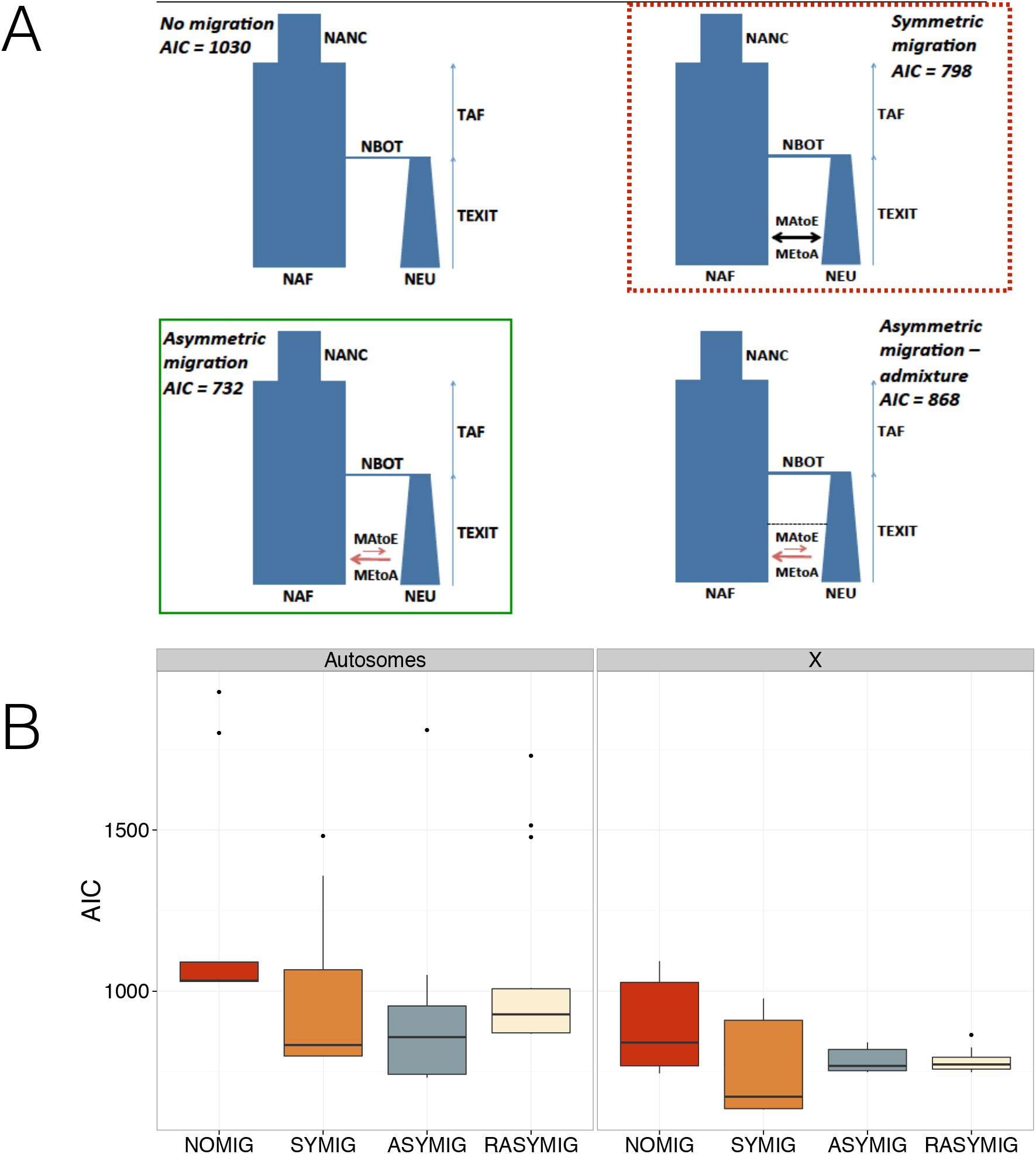
Results of the model choice analyses. A) The four demographic models tested in this study. Lowest AIC out of 10 replicates are reported for each model. The green box with a continuous line indicates the best model for the autosomal data. The red box with the dotted line indicates the best model for the X-linked data. B) Distribution of AIC for each for the autosomal and X-linked datasets across 10 replicates. Lower values of the AIC statistic indicate a better fit between the observed data and the demographic models.

**Table 2.** Demographic estimates from this study compared to the demographic estimates obtained by Laurent et al. (2011) for the same populations. For the *dadi* estimates we report the maximum likelihood estimates and the confidence interval obtained with parametric bootstrapping. The estimates from Laurent et al. (2011) correspond to the mode and the 2.5^th^ and 97.5^th^ quantiles of the posterior distribution.

| Parameters | Autosomes |  | X |  |
| --- | --- | --- | --- | --- |
|  | This study (ML) | Laurent et al. (2011)<br>ABC | This study (ML) | Laurent et al. (2011)<br>ABC |
| Current African population size ($N_{AF}$) | 4,639,014<br>(4,000,067; 5,217,550) | 3,134,891<br>(1,371,066; 28,013,950) | 7,537,910<br>(6,425,986; 9,845,444) | 4,786,360<br>(2,040,701; 29,208,295) |
| Current European population size ($N_{EU}$) | 957,941<br>(591,256; 1,917,823) | 878,506<br>(383,361; 4,775,964) | 529,902<br>(355,231; 1,146,428) | 1,632,505<br>(780,907; 4,870,580) |
| European Bottleneck population size ($N_{BOT}$) | 112,191<br>(71,831; 242,384) | 32,128<br>(15,968; 95,162) | 41,507<br>(22,529; 81,830) | 22,066<br>(14,338; 81,102) |
| African-European divergence time ($T_{SPLIT}$) | 43,540<br>(33,154; 74,498) | 12,843<br>(7,095; 31,773) | 25,999<br>(18,810; 37,809) | 16,849<br>(9,392; 33,452) |
| African expansion time ($T_{EXP}$) | 61,334<br>(24,011; 119,173) | 37,323<br>(3,636; 379,212) | 79,776<br>(44,008; 117,303) | 25,553<br>(1,698; 376,730) |
| Ancestral African population size ($N_{ANC}$) | 2,058,317<br>(1,949,391; 2,145,813) | 1,705,328<br>(609,393; 2,458,653) | 2,147,406<br>(2,040,027; 2,253,918) | 1,837,229<br>(931,637; 2,530,609) |
| Migration rate Europe to Africa ($M_{SZ}$) | 2.24<br>(1.26; 2.67) | not estimated | 1.23<br>(0.83; 1.58) | not estimated |
| Migration rate Africa to Europe ($M_{ZS}$) | 0.53<br>(0.09; 1.31) | not estimated | 1.23<br>(0.83; 1.58) | not estimated |

It has been shown that large contiguous sequence information from a sample of size two contains information about historical changes in coalescence rates (Li & Durbin, 2011). In theory, this approach should complement classical model-based inference procedures like the one presented in Figure 2, because no prior assumptions are required on how often the coalescence rate can change during the history of the sample. We therefore compared the inverse instantaneous coalescence rates (IICR, see Materials and Methods) simulated under our best neutral model (ASYMIG, autosomal data, Figure 2, Table 2) with the IICR measured with *MSMC* using the complete sequences of chromosomes 2R, 3L, and 3R (Figure 3). The simulated IICR for the three possible types of samples of size two (i.e. Sweden-Sweden, Sweden-Zambia, and Zambia-Zambia) varied according to the major demographic events implemented in the model ASYMIG: the ancestral expansion, the population split, and the population size bottleneck in the Swedish population (Figure 3B). As expected in an isolation-with-migration model, the IICR did not match exactly the effective population sizes in the model (Chikhi et al., 2018). Indeed, the most recent IICR for the Swedish and Zambian samples are slightly higher than the real effective population sizes defined by our model (Figure 3B). Also, we note that the IICR of the Zambian population after the split is lower than the Zambian effective population size in the model. This is likely caused by the increased coalescence rate of African lineages in the European population during the time when the European population has a very low effective population size. Interestingly, the IICR estimated by *MSMC* (n=2) differed from the simulated neutral IICR in many respects (Figure 3B, Figure S6). First, we observed a decrease in IICR in both populations between 100 and 1000kyrs that is not predicted by our model. Second, we did not observe any change in the MSMC-IICR values at the time of the African expansion nor at the time of the population split. Instead, around this time, we observed a steady decrease in MSMC-IICR for the Swedish population while the African population remained relatively constant. Finally, we observed a decrease in African MSMC-IICR in recent times that, again, was not predicted by our model.

**Figure 3:**
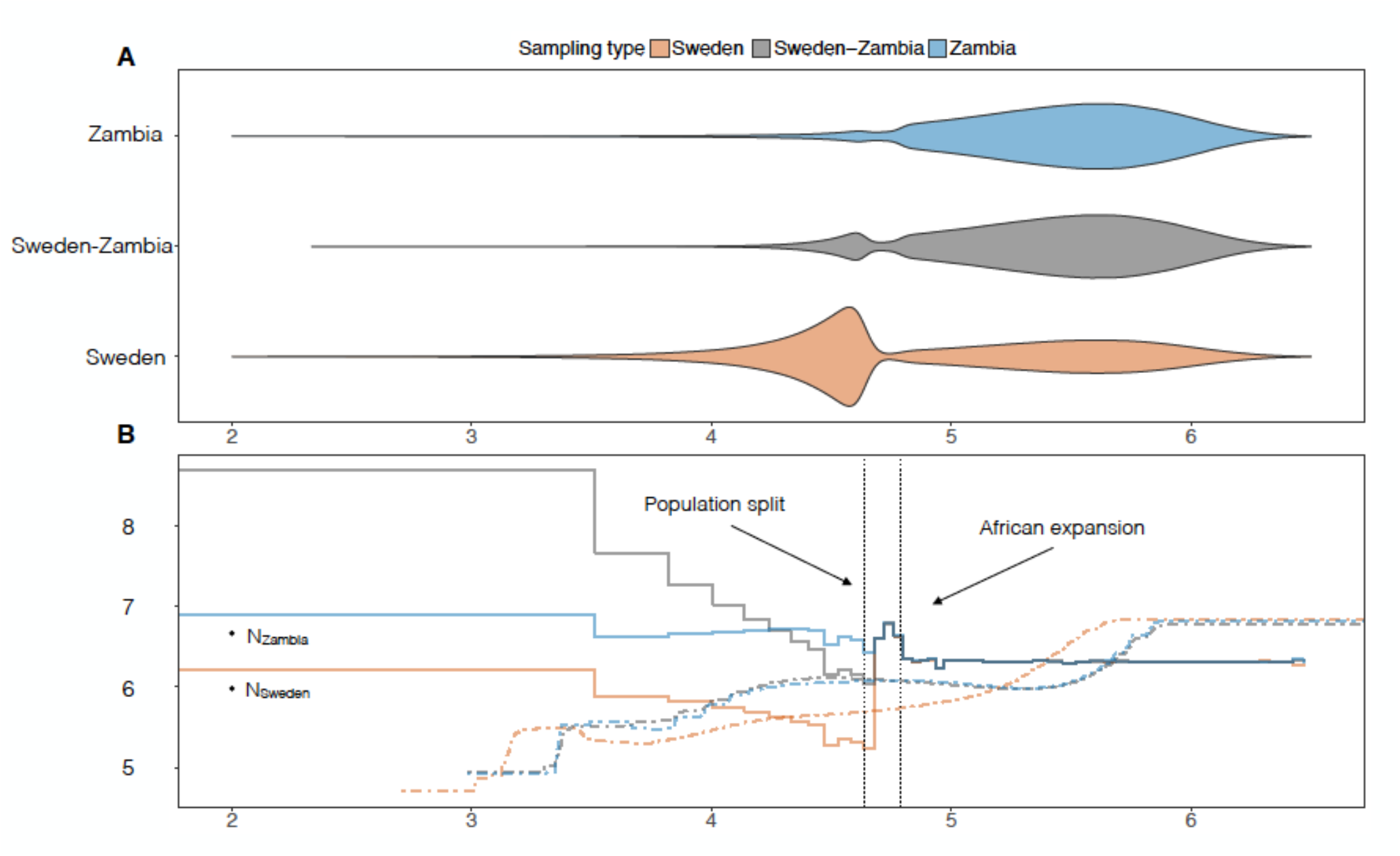
A) Distribution of coalescence times for samples of size 2 under our best demographic models inferred with *dadi* (autosomal data, Figure 2). One million simulations were conducted with the software ms using the following command line: *ms 2 1000000-T -L -I 2 2 0 -m 1 2 10.0815052248 -m 2 1 2.41228728 -n 2 0.2064966823 -g 2 91.3987189817 -ej 0.0234640378 2 1 -en 0.03305336 1 0.4436970874*. B) Comparison of the inverse instantaneous coalescence rates (IICR, solid lines) predicted under our best model (same as panel A) with *MSMC* results (dotted lines

## Discussion

The PCA results presented in Figure 1a illustrate that chromosomal inversions can have a strong impact on neutral polymorphism data. Importantly, the structure created by *In*(*2L*)*t* in the data extends several megabases beyond the inversion’s breakpoint (Corbett-Detig & Hartl, 2012; Huang et al., 2014). Therefore, we excluded all of chromosome 2L for the demographic analyses in this study and recommend that future demographic studies of natural populations of *Drosophila melanogaster* address the potential effect of this inversion prior to model fitting. Alternatively, coalescent models that explicitly account for the effect of chromosomal inversions (Guerrero, Rousset, & Kirkpatrick, 2012; Peischl, Koch, Guerrero, & Kirkpatrick, 2013) could be used to jointly take into account demographic processes and the specific patterns of recombination caused by the inversion. We note that such models could in principle be used to investigate why the genetic differentiation between African and European populations is lower for *In*(*2L*)*t* compared to the standard arrangement (Figure 1a). We speculate that lines carrying *In*(*2L*)*t* may have colonized Europe more recently than lines carrying the standard arrangement, leaving less time for drift to increase differentiation.

Our model-based demographic analyses (Figure 2, Table 2, Figure S5) confirmed that European populations do not exhibit patterns of African admixture comparable to the ones that have been measured in American and Australian populations (Bergland et al., 2016; Caracristi & Schlotterer, 2003; Duchen et al., 2013). This indicates that European natural populations of *D. melanogaster*, and ancient populations in general may be better suited for studying local adaptation at the genetic level because neutral models serving as a null hypothesis for selection detection methods do not have to account for the additional complexity caused by genetic admixture (but see Lohmueller, Bustamante, & Clarck, 2011). The new Swedish panel presented in this study therefore represents an appropriate sample to address the long lasting issue of the respective contributions of hard versus soft selective sweeps in the adaptation of *D. melanogaster* to northern latitudes (Garud, Messer, Buzbas, & Petrov, 2015; Jensen, 2014).

Nevertheless, accounting for ongoing gene flow between Africa and Europe improved the fit to the data compared to models that do not allow for migration (Figure 2b). As predicted by Li and Stephan (2006), allowing for gene exchange between Africa and Europe in the demographic model provides an older estimate for the age of the split between the two populations (Table 2, Table S2). The age of the divergence obtained from neutral autosomal under our best model (43,540 years) suggests that the split between African and European ancestral lineages occurred during the last glacial period. Another interesting consequence of including migration is that estimates for N_BOT_ (the size of the European population directly after the split) are roughly two times larger than in models without migration (Table S2). The potential effect of a less severe bottleneck and gene flow on the performance of selection detection in *D. melanogaster* remains to be investigated and is beyond the scope of this study. Interestingly, our estimation of the migration rate from Europe to Africa is lower on the autosomes than the X-chromosome (2.24 and 1.23, respectively; Table 2). This result is consistent with previous results based on microsatellite variation that showed that European admixture in urban African populations was significantly higher in autosomes compared to the X chromosome (Kauer, Dieringer, & Schlotterer, 2003).

A possible reason for the striking differences that we observed between the IICR inferred by the *MSMC* and the IICR predicted under our best neutral demographic model is the use of different genetic datasets in both cases. While our demographic model has been calibrated with genetic variation measured in short introns and third codon positions (i.e. putatively neutral loci), *MSMC* used the complete sequences of chromosome 2R, 3L, and 3R. Therefore, the proportion of loci subject to purifying selection is expected to be much larger in the *MSMC* analysis and could explain that MSMC-IICR are lower than neutral expectations for most of the populations history (Figure 3B). Another important consideration is that *MSMC* uses both the distribution of coalescent times and the length of non-recombining genetic intervals to estimate IICR values, while *dadi* only uses the former. It is possible that selective processes like selective sweeps or local distortions of the recombination rates caused by chromosomal inversions affect *MSMC* results more strongly than demographic models calibrated with the site frequency spectrum only. Finally, it cannot be excluded that *MSMC* provides a more detailed description of the true change in population sizes in European and African populations of *D. melanogaster* because it does not require restrictive arbitrary assumptions about past changes in population size. It remains to be investigated whether a continuous spatial range expansion from sub-Saharan African to northern European latitudes would predict a decreasing IICR curves such as the one inferred by *MSMC* for the Swedish sample (Figure 3B) and therefore potentially represents a more appropriate neutral model for the origin of cosmopolitan populations.

In this study, we showed that the time of divergence between the African and European population is more than three times older than previously published estimates. While previous results concluded that this split occurred at the end of the last glacial maximum (LGM), our new estimates sets the divergence time before the LGM, at a time when climatic conditions in Europe were not suitable for *D. melanogaster*. Hence, most of the neutral differentiation between current European and sub-Saharan African flies occurred while both populations were still located on the African continent. As a consequence, signatures of selection identified in European samples may potentially reflect molecular adaptations to African rather than European environments.

## Acknowledgements

We thank Roman Arguello for helpful discussions and for providing the coordinates of short introns and fourfold degenerate coding sites for the neutral set of loci. We thank Jeffrey Jensen and Thomas Flatt for helpful discussions and advices. Adamandia Kapopoulou was supported by a SNF grant to Jeffrey Jensen. Martin Kapun was supported by SNF grants PP00P3_133641and PP00P3_165836 to Thomas Flatt. Sequencing of the Swedish lines was supported by grant STE325/12-2 of the DFG Research Unit 1078 to Wolfgang Stephan.

## Auhor contributions

R.W and S.L collected biological material, A.K conducted the demographic inference with dadi. M.K conducted all bioinformatics analyses to process the data. B.P. conducted the *MSMC* analyses. S.L. conducted the IICR analyses. P.P., W.S, and S.L designed the study and wrote the paper.

